# Green algal symbionts are stably retained and provisioned in the dark despite a clear physiological cost

**DOI:** 10.64898/2026.08.17.745325

**Authors:** Dongseok Kim, Betsy Varghese, Sergio A. Muñoz-Gómez

**Affiliations:** Department of Biological Sciences, Purdue University, West Lafayette, IN, USA; Center for Plant Biology, Purdue University, West Lafayette, IN, USA

**Author notes:** Correspondence to: Sergio A. Muñoz-Gómez, Department of Biological Sciences, Purdue University, 915 Mitch Daniels Blvd., West Lafayette, IN 47907, USA.

**Keywords:** *P. bursaria*, photosymbiosis, mutualism, exploitation, parasitism, physiology, symbiont load, burden, cost, benefit, proteomics, systems biology

## Abstract

Photosymbioses—associations between heterotrophs and photoautotrophs—are widespread and indispensable in today’s ecosystems. The chloroplasts of algae and land plants, which are at the heart of most of earth’s primary production, stem from ancient photosymbioses. Photosymbioses often combine heterotrophy and autotrophy and must thus efficiently allocate resources between these two costly cellular processes. We currently lack a clear picture of how photosymbioses allocate their valuable cellular resources in response to environmental change. In this study, we combine growth assays, automated fluorescence microscopy, transmission electron microscopy, and mass spectrometry-based proteomics to explore the physiology and cellular resource allocation of the ciliate-green alga photosymbiosis of *Paramecium bursaria*. In nutrient-rich environments that resemble *P. bursaria*’s natural habitat, the maximum growth rate attained saturates regardless of light intensity. The green algae thus do not provide a benefit in nutrient-replete conditions, and the photosymbiosis primarily functions heterotrophically. The green algae occupy a remarkably similar and constant volume fractions across contrasting light environments despite displaying clear photo-physiological adaptation. This is true regardless of a clear physiological cost of the photosymbionts—aposymbiotic hosts always display higher growth rates in the dark. The host does not decrease ‘symbiont load’ in environments where green algae are not beneficial. Moreover, in the dark, the green algae are fully dependent on their hosts and take up a larger proteome mass fraction that increases with prey abundance. Differential protein expression analyses suggest that acetate and amino acids are the preferred sources of carbon and nitrogen for the green algae in the dark. The stable persistence and higher resource uptake by the photosymbionts in the dark argue against a view where hosts have full control over and selfishly exploit their symbionts.

## Introduction

Eukaryotes first evolved and diversified as heterotrophs [1]. Some eukaryotes subsequently became phototrophic by transforming an endosymbiotic cyanobacterium or eukaryotic alga into a chloroplast [2, 3]. An alternative way to become photosynthetic was for some eukaryotes to temporarily retain the chloroplasts of algal prey or establish symbiotic relationships with algae [4, 5]. The latter strategy—photosymbiosis—led to widespread and ecologically important groups such as corals and lichens [6, 7]. In shallow marine waters, for example, coral-alga photosymbioses provide a habitat for at least 25% of all marine species [8]. While many eukaryotes specialized as strict phototrophs, photosymbioses often combine heterotrophy with phototrophy—a strategy called mixotrophy [9]. The simultaneous expression of both heterotrophy and phototrophy, however, can be costly and be outcompeted by strict heterotrophs or phototrophs in many environments [10, 11]. The optimal allocation of resources between heterotrophy and phototrophy by mixotrophic photosymbioses is thus key to explaining their function and success across diverse ecosystems.

*Paramecium bursaria* offers several experimental advantages over other photosymbioses. Unlike corals and *Hydra viridissima*, the green algal photosymbionts in *P. bursaria* are vertically inherited from mother to daughter cells. This makes of *P. bursaria* an ideal system to understand nascent (early-stage) symbioses [12]. In addition, *P. bursaria* grows fast with doubling time of as short as seven hours, can be cultured axenically, and there are methods available for silencing host genes through RNA interference [13]. Numerous studies have investigated the physiological response of *P. bursaria* to different nutrient environments [14–19]. However, little is known about how, and if, the photosymbiosis optimizes resource allocation to maximize physiological performance or fitness in each environment. A recent study used volume microscopy to quantify resource investment into large organelles of individual symbiotic algae in *P. bursaria* in a single environment [20]. However, a global and highly resolved view of resource allocation in the photosymbiosis (e.g., between host and intracellular photosymbiont population) as a function of environmental change is still missing.

There has been a recognition that the nature of the relationship between symbiotic partners is not static and changes according to the environmental conditions [21–23]. However, several studies have favored the view that the host exploits its green algal symbionts in the photosymbiosis of *P. bursaria*. Lowe et al., (2016) provide experimental data to argue that the host adjusts the allocation of resources to photosymbionts (‘symbiont load’) to maximize benefit and minimize costs of carrying green algae [19]. The expansion of cell volume and increased allocation to photosynthetic machinery in symbiotic green algae compared to free-living relatives has been interpreted as the host exploiting its photosymbionts and forcing them to disproportionally invest into photosynthesis [20]. The allocation of cellular resources in response to the environment can be more precisely quantified by focusing on the proteome—proteins constitute most of a cell’s biomass and their synthesis consumes most of a cell’s energy [24–26]. Exploring how a photosymbiosis allocates its proteome resources across environments promises to offer valuable insights into how the interests of the symbiotic partners may shift as a function of environmental conditions.

Here, we investigated how host and photosymbionts in *P. bursaria* physiologically respond to each other and reallocate resources under contrasting environments. We combined growth assays, photophysiology, fluorescence and electron microscopy, and mass spectrometry-based proteomics. We showed that the photosymbionts (*Micractinium conductrix*) do not provide a measurable benefit as long as the photosymbiosis remains in a nutrient-rich environment, such as the stagnant and organic matter-rich freshwater sediments that *P. bursaria* inhabits. The green algal population, however, responds to the contrasting light environments by adjusting its cell volume, numbers, and chlorophyll contents. Despite this, the volume fraction taken by the green algae—a proxy for cost—remains relatively constant and similar across environments along the growth curve. Moreover, the proteome data suggests that the green algae take up a larger amount of resources and rely on host amino acids and acetate in the dark. Our results suggest a more active role for the green algae in photosymbiosis that can be viewed as an environment-permissible exploitation of the host by its photosymbionts.

## Results and Discussion

### Section 1: Heterotrophic metabolism dominates early and masks photosymbiont benefits in nutrient-rich environments

Unlike most marine photosymbioses like corals and radiolarians that thrive in oligotrophic (nutrient-poor) environments [7, 27], *P. bursaria* and its green algal symbionts inhabit stagnant freshwater ponds that are, comparatively, nutrient-rich or eutrophic [28]. Within this environment, *P. bursaria* has been observed to occupy the oxic-anoxic boundary close to sediments where light levels are low [29]. To investigate the physiological response of the photosymbiosis of *P. bursaria* to contrasting environments, we first performed growth experiments in nine combinations of irradiance and prey bacterial concentrations. We aimed to recreate well differentiated environments that reflected the natural habitat of *P. bursaria*. To do this, our culture medium was based on a wheat grass infusion whose composition has been calculated to contain ∼625 mg/L of protein and ∼850 mg/L of carbohydrate, among other nutrients [30]. This medium thus provided a considerable amount of dissolved nutrients and ensured robust and reproducible growth across conditions. Similar plant-infused media have previously been shown to produce differences in the growth rate and yield of *P. bursaria* [19, 31]. We developed a workflow for the (semi-)automated counting and size estimation of *P. bursaria* cells using a wide-field fluorescence microscope with a motorized stage. This allowed us to obtain highly resolved growth curves and reliable estimates of cell sizes along the growth curve. Together, these data provide a much more precise estimation of growth physiological parameters compared to previous studies.

In nutrient-rich environments, that may more closely resemble the natural habitat of *P. bursaria*, there are no differences in maximum growth rate (*μ_max_*) across environments—growth rate saturated at a value of ∼0.4 day^-1^ regardless of irradiance and prey bacteria concentration (**Fig. 1A**; **Fig. 1B**). Similarly, the maximum cell sizes (*V_max_*) achieved did not vary significantly across environments. Both *μ_max_* and *V_max_* are consistently reached at about day two (**Fig. 1B**; **Fig. S1**). This suggests that the photosymbionts do not provide a measurable physiological benefit to their hosts for at least the first 4 days of growth (or about two generations) or as long as nutrients remain in excess (**Fig. 1A**). The unresponsiveness of growth rate to environmental changes, as well as the small effect of different bacterial prey concentrations, is arguably caused by a saturating amount of dissolved nutrients that support the *μ_max_* that the photosymbiosis can achieve.

**Figure 1.**
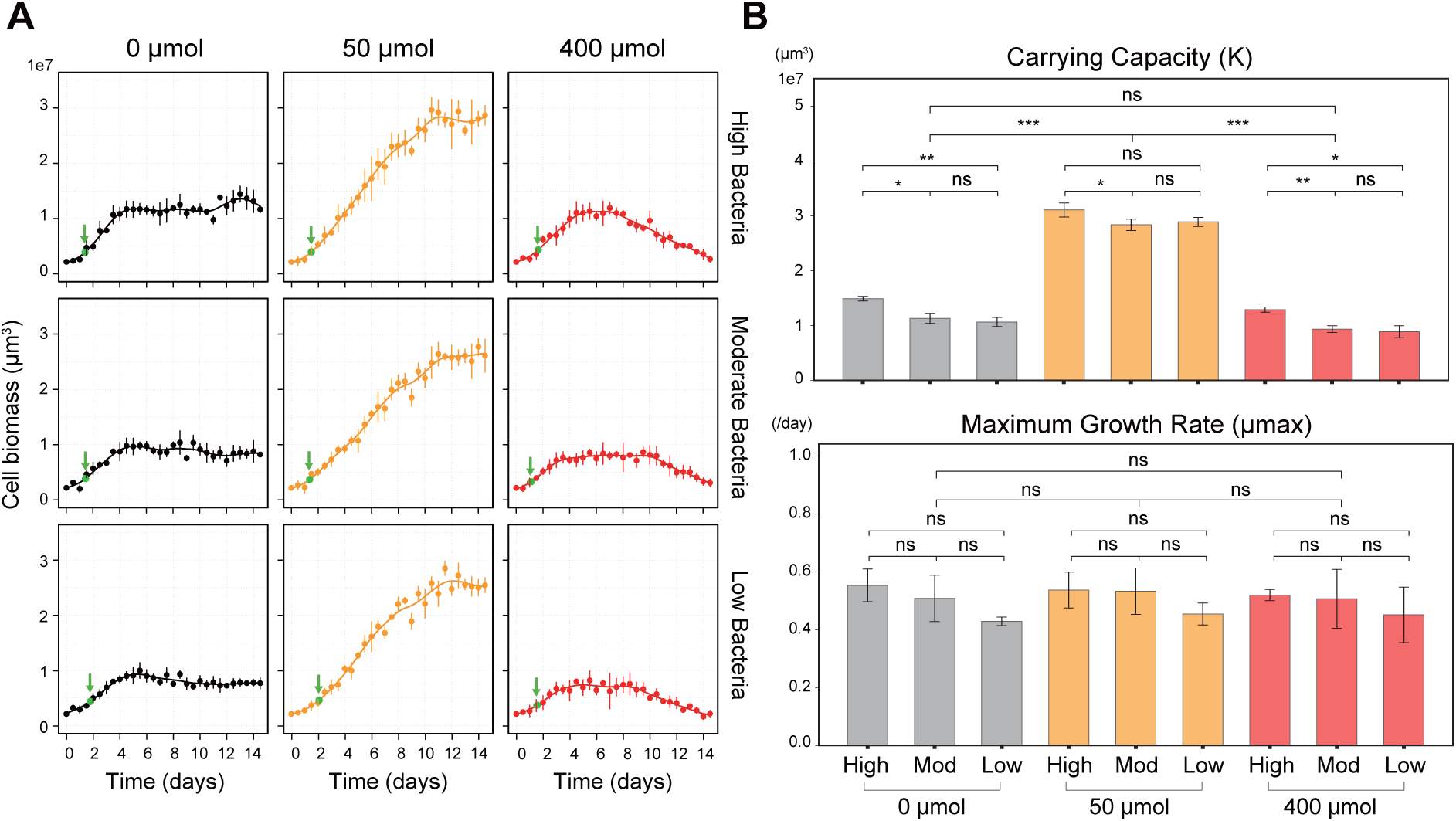
Global growth physiology of the *P. bursaria* holosymbiosis and its green algal photosymbionts in nutrient-rich environments. (A) Biomass-based growth curves of the holosymbiosis under nine combinations of irradiance (0, 50, and 400 µmol·m^-2^·s^-1^) and three prey bacteria concentration [low (6.4 × 10^5^ cells mL ¹), moderate (1.28 × 10^8^ cells mL ¹), and high (6.4 × 10^8^ cells mL ¹)]. The biomass of the holosymbiosis is calculated as the product of cell number and volume. The point of maximum growth rate on each curve is marked with a green arrow. **(B)** Estimated growth parameters under each of the nine conditions tested. The upper panel shows the maximum growth rate, and the lower panel shows the maximum population biomass that the environment can sustain. Significance of pairwise differences was assessed by t-test, with results shown above each bar (ns, not significant; *, *P* < 0.05; **, *P* < 0.01; ***, *P* < 0.001).

Irradiance exerts a much stronger influence on growth dynamics than prey bacterial concentration (**Fig. 1A**). In a moderate light environment (50 µmol photons·m^-2^·s^-1^), the photosymbiosis was able to sustain growth for much longer (∼12 days) which resulted in a much higher maximum population size (and total biomass) compared to other environments—the host benefits the most from the photosymbionts under an optimal light environment after nutrients have been sufficiently depleted. In the dark, the maximum population biomass achieved is about three times lower than that achieved under a moderate light intensity. The lack of a host benefit from non-photosynthesizing green algae thus manifests as a reduced maximum population size after most nutrients have been depleted at stationary phase—growth is sustained for a shorter time (**Fig. 1A**). There is an additional cost seen as a death phase or population size decline that appears after day ∼8 10 under the high irradiance (**Fig. 1A**). This is presumably due to cellular damage by reactive oxygen species (ROS) ultimately stemming from excess light [32]. The integrated biomass accumulated across time is thus much lower under high irradiance than in the dark (**Fig. 1A**).

An excess of nutrients maintained by daily culture passaging has been shown to maintain the same *μ_max_* in dark and light environments [33]. We confirmed this result here across a broader range of environments (**Fig. 1**). This suggests that a steady supply of nutrients suppresses the benefits conferred by photosynthesizing (i.e., cooperating) green algae. The photosymbiosis thus must prioritize a heterotrophic physiology when nutrients are in excess. As nutrients are depleted during the growth curve, the physiology of the photosymbiosis progressively switches to, or increasingly prioritizes, photoautotrophy. This agrees with the view that photosymbioses provide an advantage over heterotrophic relatives as nutrients become scarce or in oligotrophic conditions [27, 34]. In the photosymbiosis of *P. bursaria*, however, a stronger reliance on heterotrophy may be an adaptation to its more nutrient-rich habitat in stagnant freshwater ponds.

### Section 2: Resource allocation to photosymbionts stays approximately constant across contrasting light environments despite substantial photo-physiological adaptation

At the broadest level, one expects *P. bursaria* to control the allocation of resources to its intracellular green algal population. It has been reported, for example, that the total number of green algae per host cell varies with irradiance [19]. The amount of biomass (or volume) taken up by the green algal population serves as a proxy for resources invested into phototrophy over heterotrophy. To investigate coarse resource allocation in a photosymbiosis, we followed the dynamics of the intracellular green algae in *P. bursaria* along the growth curve. We counted and measured the sizes of hosts and photosymbionts from several *P. bursaria* cells using tomograms obtained with a confocal fluorescent microscope.

These experiments revealed opposite trends in the number and size of the green algae between dark and light environments. Green algal size increases under light, being ∼2 4-fold larger than in the dark (**Fig. 2A**); this creates a higher surface area-to-volume ratio for the green algae in the dark (**Fig. S2**). On the other hand, the number of green algae is ∼2 2.5-fold higher in the dark than in the two light environments (**Fig. 2B**). These trends are largely unaffected by the concentration of prey bacteria (**Fig. S3**). The green algae photo-acclimated as seen by different chlorophyll amounts despite following similar size-number dynamics at moderate and high irradiance—per-alga chlorophyll contents decrease rapidly in the dark and high-light environments (**Fig. S4**). Maximum photochemical efficiency of PSII (Fv/Fm) and non-photochemical quenching (NPQ) also showed changes across light environments and along the growth curve (**Fig. S4**). An increase in green algal size in the light is arguably due to photosynthate accumulation and cellular growth under arrested division imposed by the host (by, for example, controlling the supply of a key nutrient). We quantified the fractional amount of starch across light environments using transmission electron micrographs and found that chloroplast starch granules occupy ∼2-6% of the green algal cells (**Fig. 2C**; **Fig. S5**). On the other hand, the increase in the number of green algae (and decrease in size) is due to an initial acceleration of cell division in response to dark exposure [35, 36] without subsequent photosynthate accumulation (**Fig. 2B**). The division rate of the green algae later slows down to match the host’s growth rate, and a newly stable larger intracellular green algal population size is achieved; green algae appear to slowly decline in number toward the end of the 14-day growth curve (**Fig. 2B**).

**Figure 2.**
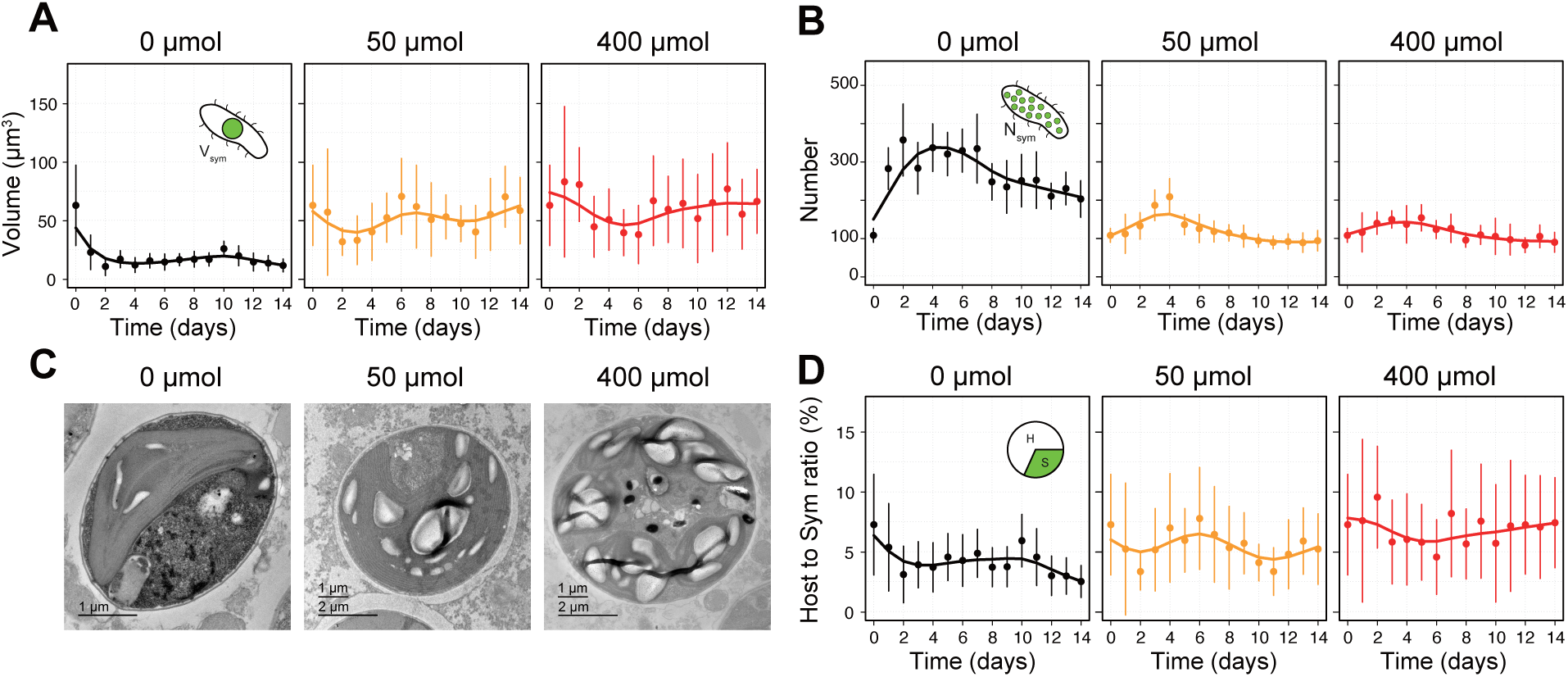
Photosymbiont dynamics across time and nutrient environments. **(A)** Volume of a single photosymbiont cell along the growth curve. Cell volume is calculated using the measured cell diameter by assuming a perfect sphere. **(B)** Total number of photosymbionts within a single host cell along the growth curve. **(C)** Representative TEM micrographs of photosymbionts under the three light conditions in the high-bacteria condition. **(D)** Proportion of host volume occupied by total photosymbiont biomass (number × volume) along the growth curve. In (A), (B), and (D), data for only the highest prey bacteria concentration are shown. Other prey bacteria concentrations are shown in Fig. S3.

Salsbery and DeLong (2021) observed a similar opposite trend between green algal size and number but in response to cold and hot temperatures (20 and 32°C). The size of the green algae was larger, and their number lower, at the colder temperature of 20°C. The opposite was observed at 32°C [37]. The larger size and lower number of the green algae coincided with a growth rate about twice as high in the lower temperature [37]. It is possible that higher temperatures decrease the efficiency of photosynthesis making the green algae less beneficial to their hosts and thus reducing the fitness of the photosymbiosis. Altogether, these observations suggest that a smaller population of larger-sized green algae may be more favorable or even optimal for the photosymbiosis.

The opposite trends in green algal number and size compensate each other and produced a near-constant photosymbiont-host volume ratio (here and in Salsbery and DeLong (2021); **Fig. 2D**). This can also be seen in how the total green algal biomass tracks that of the hosts along the growth curves (**Fig. 1A**; **Fig. S6**). The photosymbiont-host volume ratio varies within a narrow range of ∼2-10% and is slightly larger in light environments (**Fig. S3**), although the differences are not significant. These results suggest that the material costs of the green algal population— measured as a volume fraction proxy—are approximately fixed across environments and do not change considerably even as nutrients are depleted along the growth curve.

### Section 3: Green algae impose a cost and derive a much greater benefit in the dark

The near-constant photosymbiont volume fraction reported here conflicts with the view that the host exploits its green algal symbionts and readily adjusts their load to maximize fitness across environments [19]. Such a view predicts that ‘symbiont load’ (or host resources devoted to symbionts) is lowest in environments where the photosymbionts do not cooperate or contribute to overall fitness (i.e., dark). In agreement with our findings, previous studies have shown that the green algal population can persist in the dark for several weeks [14, 38, 39]—gradual decline and eventual loss occur under starvation in nutrient-poor environments [40, 41]. It may thus be argued that under nutrient-rich environments, the green algae are stably maintained because they represent little or no cost to their host. To test this directly, we cultured the holosymbiosis (ciliates associated with green algae) under a one-time- and regular-feeding regimes. In addition, we created aposymbiotic hosts by curing the holosymbiosis of their green algae (cycloheximide treatment in the light; see **Materials and Methods**) and evaluated their growth dynamics in the dark. The growth curves revealed that a regular supply of nutrients maintains a larger green algal population (**Fig. 3A**), and that aposymbiotic hosts always grow faster than holosymbiotic hosts in the dark (**Fig. 3B**). This suggests that the green algae pose a significant cost to their host that could, in principle, be reduced (e.g., by means of digestion or expulsion) in a shorter time frame to improve fitness. Costly photosymbionts are thus maintained at a constant load despite posing a clear physiological cost as long as there is a constant supply of nutrients.

**Figure 3.**
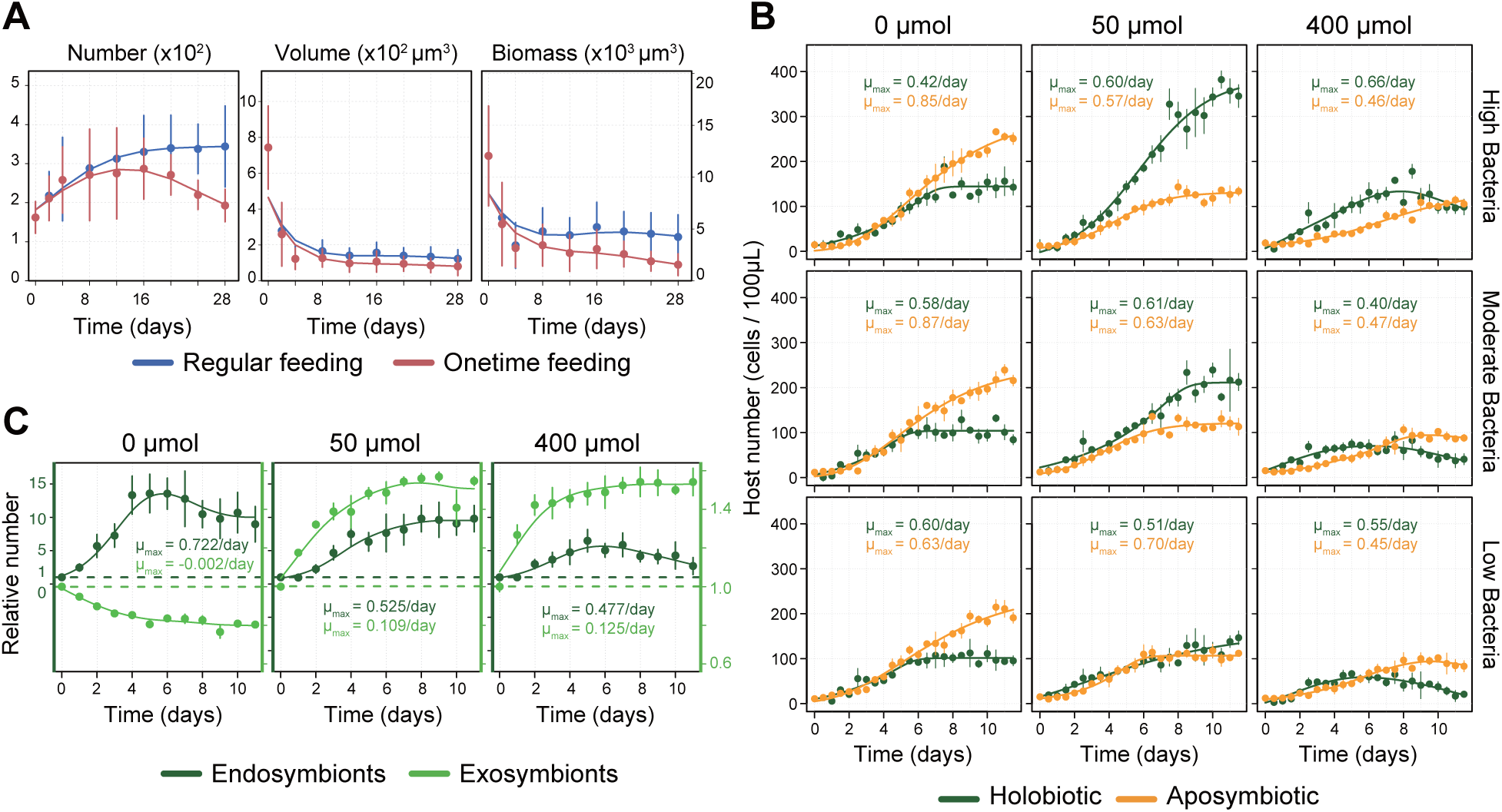
Costs to the host and benefits to the photosymbiont. **(A)** Changes in the number, volume, and biomass (number × volume) of symbionts within the host under two different feeding regimes: regular feeding, in which fresh medium and prey bacteria are supplied every 4 days, and one-time feeding, in which no resupply is provided. These experiments were performed in the dark and a high prey bacteria concentration. **(B)** Growth curves of holosymbiotic versus aposymbiotic *P. bursaria* across the nine conditions. **(C)** Growth curves of endosymbiotic and exosymbiotic green algae. The y-axis is normalized OD880 relative to the initial OD880 value. The left y-axis corresponds to endosymbionts whereas the right y-axis to exosymbionts. Note different scales.

The above may suggest that the photosymbiosis has evolved to nourish and maintain the green algae in the dark. This can be viewed as an adaptation to both short-term diel (i.e., night) and long-term seasonal (i.e., winter) darkness. To test whether the green algae derive a benefit from their hosts in the dark, we isolated and successfully cultured *Micractinium conductrix*, the green algal symbionts of *P. bursaria*, in wheat grass infusion medium (see **Materials and Methods**). Growth curves across light environments revealed that the exosymbiotic algae cannot grow in the dark when outside their hosts (**Fig. 3C**). This suggests that the host provides a uniquely nutritious environment to their photosymbionts—the green algae not only survive but also grow and multiply when inside their hosts in the dark. Dark growth is thus a major advantage that green algae derive from engaging in photosymbiosis—an external environment is unlikely to offer the right amount and combination of nutrients to support substantial heterotrophic growth of exosymbiotic algae. Different degrees of inter-dependence between symbiotic partners have been reported across *P. bursaria* species [31]. We suggest that *P. bursaria* strain CCAP 1660/18 (recently renamed as *Paramecium deuterobursaria* [28]) has evolved the ability to heterotrophically maintain its intracellular green algae for a longer time or, alternatively, the green algae have an increased ability to sustain themselves heterotrophically.

### Section 4: Proteome investment into photosymbionts in the dark is greater despite a diminished investment into photosynthesis

Investigating cellular resource allocation across environments reveals what functions and organelles the photosymbiosis prioritizes to maximize growth and survival. In *P. bursaria*, this has so far been studied using light microscopy or volume electron microscopy under a constant environment [20]. These approaches, however, only provide a very coarse view of cellular resource allocation in the photosymbiosis. The proteome offers a more direct proxy for material costs and a better resource to investigate both coarse- and fine-grained cellular resource allocation.

We employed label-free quantitative proteomics across our nine nutrient environments sampled after photo-acclimation had occurred, enough biomass had accumulated, and at least four generations had elapsed. In agreement with growth physiology and volume-based cellular allocation (**Fig. 1**; **Fig. 2**), the proteome response is mostly driven by irradiance (**Fig. 4A**). We estimated the proteome fractions taken by host and photosymbionts. The data show that the intracellular green algal population occupy a relatively narrow proteome fraction of 8.0 14.3% (**Fig. 4B**). These values are similar to the photosymbiont volume fractions estimated by microscopy (**Fig. 2D**). However, the photosymbiont proteome fraction is significantly larger in the dark at 11.3-14.3% compared to 8.0-9.6% in the light (**Fig. 4B**). This is contrary to expectations and is opposite to the (non-significant) trends observed for volume fractions across light environments (**Fig. 2D**). The discrepancy between proteome mass and volume fractions is partially resolved by considering that about ∼2-6% of the green algal biomass in the light is occupied by starch and lipids (**Fig. 2C** and see [20]). The material (proteome) costs of the photosymbionts increase in those conditions under which the symbionts do not offer a benefit to their hosts.

**Figure 4.**
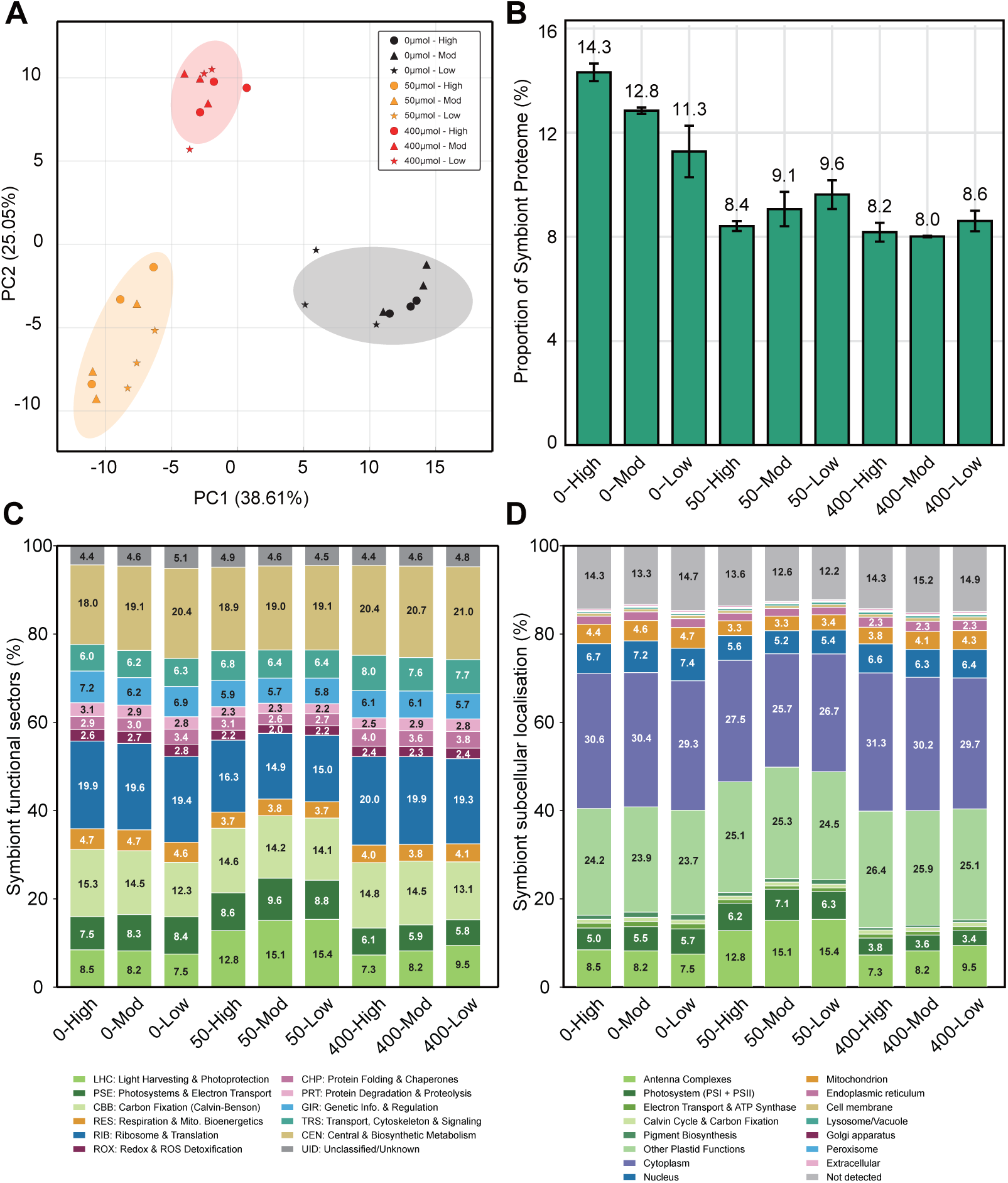
Proteome resource allocation between and within each symbiotic partner in the *P. bursaria* holosymbiosis. **(A)** PCA plot of proteome profiles of relative protein abundances across the nine conditions investigated. Each individual biological replicate is shown. **(B)** Proportion of the holosymbiosis proteome allocated to the population of the green algae for each of the nine conditions. **(C)** Proteome resource allocation in the photosymbionts across functional sectors for each of the nine conditions investigated. Symbiont proteome partitioned into eight functional sectors based on gene annotations. **(D)** Proteome resource allocation in the photosymbionts across organelles for each of the nine conditions investigated as predicted by DeepLoc v2.1. The plastid sector (shown in green) was further subdivided into six functional subsectors. See Fig. S7 for proteome resource allocation across functional sectors and organelles in the host. See Table S1 for source data.

A response to prey bacteria concentration becomes apparent in the proteome fraction allocated to the green algal population. In the light, the symbiont proteome fraction increases (or stays constant) with decreasing prey bacteria (**Fig. 4B**). As host heterotrophy becomes limiting in the light at lower prey bacteria concentrations, the investment into actively photosynthesizing green algae increases to compensate for a lower heterotrophic carbon influx. This is consistent with host control of photosymbionts in the light [19, 20]. In the dark, the opposite is true: photosymbiont proteome fraction increases from ∼11.3 to 14.3% with increasing prey bacteria (**Fig. 4B**). The green algae take up more resources as heterotrophic carbon influx increases at higher prey bacteria concentrations. The difference in growth rate between holo- and aposymbiotic strains in the dark is bigger when prey bacteria are most abundant (**Fig. 3**). This in agreement with a higher proteome cost of the photosymbionts to the host under these conditions (**Fig. 4B**). Altogether, these observations are at odds with host control and suggest uncooperative or selfish behavior on the part of the green algae.

Across light environments, the photosymbionts undergo more proteome re-allocation than the host (**Fig. 4C**; **Fig. 4D**; **Fig. S7**). This is primarily a function of investment into chloroplast and photosynthesis. The photosymbionts invest a greater amount of proteome resources, about 50%, into chloroplast and photosynthesis in moderate light than in the dark and high light (**Fig. 4C**; **Fig. 4D**). This corresponds with the highest growth yield achieved by the photosymbiosis (**Fig. 1**) and suggests that proteome resources are diverted from growth to support photosynthetic output and host nutrition under optimal light conditions [20]. A decrease to ∼40% in chloroplast investment is observed in both dark and high light. In the dark, this lower chloroplast investment is achieved by decreasing light harvesting complexes and carbon fixation machinery (**Fig. 4D**). In high light, however, this is achieved mainly by decreasing both light harvesting complexes and photosystems (**Fig. 4D**). The proteome resources released by a lower allocation to the chloroplast are primarily diverted to the cytoplasm, nucleus, and mitochondria—most of the cytoplasmic proteome is devoted to the translation machinery (e.g., ribosomes and associated proteins; **Fig. 4C**; **Fig. 4D**). These patterns thus reveal a prioritization of protein synthesis (growth) and heterotrophic metabolism (mitochondria) by the green algae in the dark. Under high light, however, a larger translation proteome sector arguably does not support growth but maintenance, i.e., protein turnover. In accordance, investment into glutathione/thiol-based peroxide detoxification (including upstream sulfur assimilation metabolism) [43], protein re-folding, and proteolysis increases to cope with oxidative photodamage (**Fig. 4C**; **Table S5**). In response to lower photosynthetic investment by the green algae in the dark, the host increases investment into central and biosynthetic metabolism and translation machinery (**Fig. S7**). In summary, a larger proteome mass fraction devoted to heterotrophically growing green algae in the dark, which themselves invests less into photosynthesis, argues for a view where the photosymbionts are profiting from, instead of serving, their hosts.

### Section 5: Green algae use host-derived acetate and amino acids as major sources of carbon and nitrogen

It is thought that in the *P. bursaria* photosymbiosis, the host provides a nitrogen source in the form of amino acids [45, 46], whereas the photosymbionts provide a carbon source in the form of maltose [47]. In the dark, however, the green algal population must be maintained heterotrophically by host phagocytosis after initial photosynthate reserves are depleted (**Fig. 2B**; **Fig. S6**). Indeed, a smaller size where surface area-volume ratio is larger, as reported here, is arguably advantageous for increasing nutrient uptake by the heterotrophically growing green algae (**Fig. S2**). Nutrient exchange between the symbiotic partners in the dark has not yet been explored. The quantitative proteome data generated here offer insights into the metabolic exchange between symbiotic partners.

Glutamine has been suggested to be transferred from host to photosymbionts [45]. In agreement with this, the ciliate host enzyme responsible for synthesizing glutamine (from glutamate and ammonia), glutamine synthetase (GS), displays an expression profile that peaks at moderate irradiance (**Fig. 5A**). A similar pattern is observed for the ciliate host ammonium uptake transporter (AMT) (**Fig. 5A**). Complementarily, the green algae express a GOGAT in the light (**Fig. 5B**) which consumes glutamine to generate glutamate using photosynthetically reduced ferredoxin—glutamate then donates its amino group for the synthesis of other amino acids. These expression patterns in the light, however, also suggest that the green algae do not prioritize glutamine as a source of organic nitrogen in the dark.

**Figure 5.**
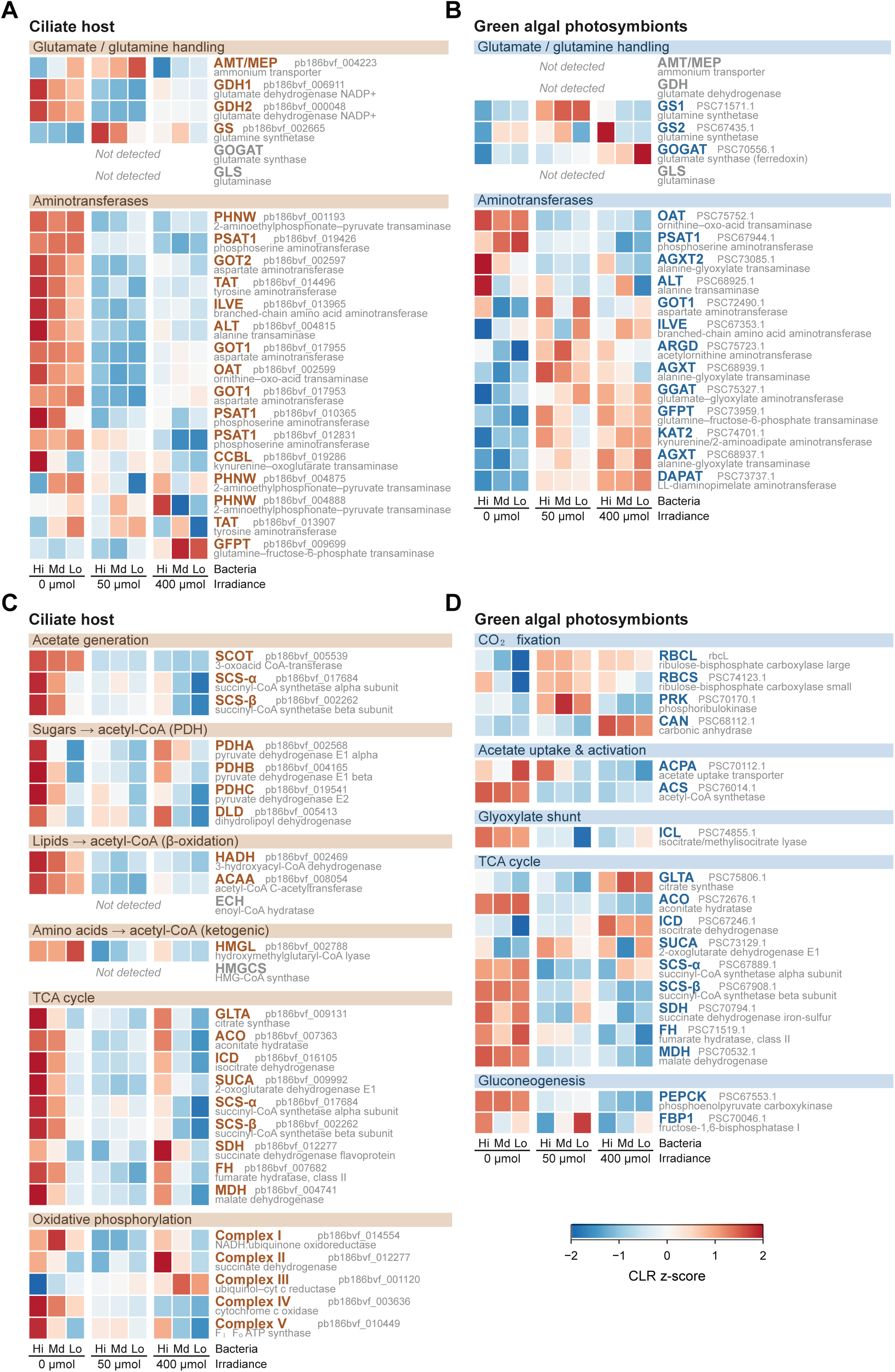
Predicted carbon and nitrogen metabolic exchange between the ciliate host and its photosymbionts. Heatmaps of CLR (centered log-ratio) z-scores of proteins involved in nitrogen and carbon exchange across the nine conditions; color indicates relative abundance from low (blue) to high (red), and “Not detected” marks proteins absent from a proteome. (A) Nitrogen metabolism in the ciliate host and (B) the photosymbionts. (C) Carbon metabolism in the ciliate host and (D) the photosymbionts.

In the dark, the ciliate host expresses a broad program for protein hydrolysis, amino acid degradation, and transamination—the typical signature of a phagotroph. This, in principle, can make all amino acids available to the green algae. Among these dark-enriched proteins in the ciliate host is glutamate dehydrogenase which catalyzes the reversible reductive amination of α-ketoglutarate into glutamate (**Fig. 5A**). The green algae overexpress a few transaminases (e.g., PSAT1, OAT, AGXT2) in the dark that use glutamate as a substrate to generate other amino acids (serine, proline, alanine) (**Fig. 5B**). Glutamate, however, cannot by itself sustain the biosynthesis of all proteinogenic amino acids. The green algae must thus import a set of several different amino acids from their host. This is consistent with previous work that showed that the green algal photosymbionts can import a broad array of amino acids from their host [46, 48]. The green algae thus appear to switch from a preference for glutamine in the light to a broader set of amino acids that derive from host phagotrophy in the dark. The photosymbionts are not simply recycling nitrogen waste from the host but consuming valuable metabolites—the host needs to invest resources to increase amino acid flux to sustain its own photosymbionts.

As a source of organic nitrogen, amino acids may serve as both nitrogen and carbon sources. However, a very high flux may be required to accomplish both roles. The green algae may thus supplement amino acid uptake with an additional carbon source. Indeed, the acetate uptake transporter (ACPA) and acetate-activating enzyme acetyl-CoA synthetase (ACS) are among the most upregulated proteins by the green algae in the dark (**Fig. 5D**). This suggests that acetate may be used by the green algae as a carbon (and also energy) source in the dark. Once activated as acetyl-CoA, acetate enters the TCA for full oxidation and energy conservation. This acetate can also provide carbon skeletons via isocitrate lyase (that bypasses TCA cycle decarboxylation) that allow for acetate carbons to be extracted as oxaloacetate from the TCA cycle via phosphoenolpyruvate carboxykinase (PEPCK) (**Fig. 5D**). On the ciliate host side, an acetate:succinate CoA-transferase (ASCT) and succinyl-CoA synthetase (SCS) enzymes create a cycle which allows for substrate-level phosphorylation while producing acetate as a byproduct (**Fig. 5C**). This cycle is fed acetyl-CoA that ultimately derives from the catabolism of prey bacteria macromolecules (sugars, proteins, and fatty acids). The ASCT-SCS cycle provides ATP alongside mitochondrial respiration, both pathways of which are enriched in the dark (**Fig. 5C**). In summary, these protein expression profiles suggest that the photosymbionts import acetate and several amino acids from their hosts as carbon and nitrogen sources to survive and grow in the dark.

## Conclusions

A commonly accepted view of photosymbiosis is that the host enslaves or exploits the algal symbionts. This view assumes strong host control of the photosymbionts and predicts that symbiont load should be minimal in environments where the photosymbionts provide no benefit. We show, however, that symbiont load or cost remains remarkably constant across drastically contrasting environments, and that the green algae are stably maintained for weeks as long as nutrients are regularly replenished. Indeed, “RNA-interference collisions” have been proposed as a mechanism evolved by the green algae to prevent host digestion across fluctuating ecological conditions [49]. Moreover, we also showed that the photosymbionts not only do not provision their hosts with photosynthate in the dark but take up more proteome resources from their hosts than they do in the light. These observations suggest an environment-permissible degree of exploitation of the host by the green algae.

## Material and Methods

### Strain information and culture maintenance

All experiments were conducted using *Paramecium bursaria* strain CCAP 1660/18. Stock cultures were maintained at 24 °C under a 12:12 h light/dark cycle with an irradiance of 50 μmol photons m ² s ¹ from 5000K cool white LED lights. The cells were grown in wheat grass medium (WGM) prepared by adding 2.5 g/L of wheat grass powder (Anthony’s organic wheatgrass powder) to ultrapure water (ThermoFisher Scientific GenPure) and boiling for 5 minutes. The solution was then filtered through Whatman Grade 1 filter paper and a 0.22 μm filter, supplemented with 0.5 g/L Na HPO, adjusted to pH 7.0, and sterilized by autoclaving at 121 °C for 30 minutes. Prior to subculturing, the medium was supplemented with filter-sterilized β-sitosterol to a final concentration of 0.8 μg/mL and inoculated with heat-killed *Enterobacter cloacae* to a final concentration of 6.4 × 10^8^ bacterial cells mL ¹). To prepare the heat-killed bacteria, *E. cloacae* was cultured in tryptone-glucose-yeast extract (TGY) medium at 28 °C for 5 hours, washed with deionized water, and heat-treated at 90 °C for 1 hour. After heating, the pellet was washed twice and resuspended in autoclaved PBS buffer. Bacterial inactivation was tested by growing aliquots on TGY agar plates overnight at 28 °C.

### Growth assays

*P. bursaria* cultures were grown under continuous illumination at 0, 50, and 400 μmol photons m□² s□¹ from 5000K cool white LED lights for 14 days. Three prey bacterial (*E. cloaceae*) concentrations [low (6.4 × 10^5^ cells mL□¹), moderate (1.28 × 10^8^ cells mL□¹), and high (6.4 × 10^8^ cells mL□¹)] were combined with the three above irradiances to create nine nutrient environments. Three biological replicates were done per condition. To quantify growth curves, samples of 100 µL were withdrawn every 12 hours and fixed with a mixture of 2.5% glutaraldehyde and stained with the fluorescent dyes Hoechst 33258 (Biotium #40044) and PKmem (Spirochrome CY-SC062). Imaging was done at 10× magnification using a Zeiss Axio Observer Z1 microscope equipped with a motorized stage and standard Zeiss filter cubes. To facilitate cell segmentation for cell counting, three fluorescence channels were employed: DAPI for DNA/nucleus staining (Ex 365, FT395, Em 445/50), DsRed for membranes (Ex 545–550/25, FT570, Em 605/70), and Cy5 for algal chlorophyll autofluorescence (Ex 640/30, FT660, Em 690/50). The acquired images were merged to generate composite images, which were then used for analysis. Cell segmentation and counting were done with Cellpose and ImageJ macro scripts. A custom Cellpose model was trained on over 30 manually annotated fluorescence images. Growth parameters were estimated using the easylinear method (fit_easylinear [50]) in the growthrates R package, with the default window (h=5, quota = 0.95). µmax was taken as the maximum log-linear slope computed per replicate (mean ± SD). Custom R scripts were used to graph growth curves.

### Establishment of aposymbiotic *P. bursaria*

To generate aposymbiotic *P. bursaria* strains, cells were washed with fresh WGM to remove debris. The cells were then transferred to a culture flask with WGM and heat-killed *E. cloacae* at a final concentration of 3.2 × 10^7^ cells mL ¹. Aposymbiosis was induced by treating the culture with cycloheximide (Thermo Scientific) at a final concentration of 10 µg/mL. The cultures were maintained at 50 μmol photons m ² s ¹ and 24 °C under continuous light for 7 days. The complete elimination of algal symbionts was confirmed by the absence of chlorophyll autofluorescence at 40× magnification under a Zeiss Axio Imager M1 microscope.

### Symbiont isolation, culture, and growth assays

To isolate green algal endosymbionts, *P. bursaria* cells maintained in WGM were washed five times with Bold’s Basal Medium (BBM) supplemented with soil extract (S/BBM), starved for 24 h in fresh S/BBM to clear ingested material, and then washed an additional five times [51]. The washed ciliates were transferred onto BBM plates (3–5 cells per plate) supplemented with beef extract (ESFI) and an antibiotic mixture of ampicillin and streptomycin at 1% and 0.25% (w/v), respectively, added as 50 μl per plate, modified from Spanner et al. [51], where they were mechanically disrupted using a sterile cell spreader to release the algal cells. The plates were incubated at 21 °C under a 12:12 h light:dark cycle, and emerging green colonies were maintained via weekly subculturing in liquid BBM + ESFI medium, with purity verified by checking for the absence of contaminating bacterial colonies on agar plates. For subsequent growth assays, the isolated exosymbiotic algae were harvested, washed, and inoculated into either WGM at a 4:15 (v/v) ratio. Triplicate cultures for each medium were maintained at 24 °C under 0, 50, and 400 μmol photons m ² s ¹. Algal growth was monitored daily for 14 days by measuring the optical density at 880 nm on an Open Colorimeter Plus (IO Rodeo) and by performing direct cell counts using a hemocytometer (Hausser Scientific) under a Motic AE2000 microscope.

### Quantification of host and symbiont volume, and symbiont load

To estimate the cellular volume of the *P. bursaria* host, we utilized the segmented micrographs generated by Cellpose for cell counting. The host cells were modeled as prolate spheroids, and their volumes were calculated based on the measured major and minor axes. We employed a custom ImageJ macro script to filter the segmented regions of interest (ROIs) and exclude debris based on the following morphological criteria: area size (800 μm^2^ < Area < 5,000 μm^2^), solidity (> 0.8), and ellipse fit ratio (0.9 < Ratio < 1.1). For the filtered ROIs, the best-fit ellipse parameters were used to calculate the volume using the standard formula for a spheroid. For the symbiotic algae, each symbiont was assumed to be a perfect sphere. The diameter of each symbiont was measured from the micrographs, and individual volumes were determined using the standard formula for the volume of a sphere. To quantify the number of symbionts per host cell, Z-stack images were acquired using a 40x objective lens in a Zeiss Axio Imager M1 microscope. The autofluorescence signal of the symbionts was detected via the Cy5 channel, and automated segmentation and counting were performed using the StarDist [52] and TrackMate [53] plugin in ImageJ, utilizing the pre-trained default model. The total symbiont volume within a host was calculated as the sum of all individual symbiont volumes.

### Transmission electron microscopy

Samples were fixed with 2.5% glutaraldehyde in 0.1 M sodium cacodylate buffer, post-fixed in 1% osmium tetroxide containing 0.8% potassium ferricyanide, and *en bloc*-stained in 1% aqueous uranyl acetate. They were then dehydrated with a graded series of ethanol and transitioned into EMbed-812 resin using acetonitrile. After transitioning to 100% resin, the resin-embedded samples were hardened in a 60°C oven. Thin sections of 80 nm in thickness were cut on a Leica EM UC6 ultramicrotome and stained with 2% uranyl acetate and lead citrate. Sample preparation was performed by staff at the Purdue Electron Microscopy Center (RRID:SCR_022687).

Transmission electron microscopy (TEM) images were acquired with an FEI Tecnai 12 TEM equipped with a Gatan ORIUS side-mount CCD camera. The microscope was run with a tungsten filament electron gun, operating at 80 kV. The photosymbiont volume was estimated by assuming a spherical shape, whereas the starch volume was calculated as that of an ellipsoid from its measured major and minor axes as measure on TEM images using Fiji.

### Measurement of photosynthetic parameters

Two methods were used to assess photosynthetic performance. First, chlorophyll fluorescence parameters were measured using a FluorCam (Photon Systems Instruments) imaging system capable of capturing chlorophyll fluorescence kinetics, including its multispectral version for detecting multiple fluorescence signals. This protocol involved a brief period of dark adaptation (15 minutes), followed by a saturating light pulse to determine maximum fluorescence parameters in dark-adapted cells, including Fo, Fm, Fv, and Fv/Fm. After dark adaptation, actinic light (400 μmol photons m ² s ¹) was applied continuously for 10 minutes, during which additional saturating pulses were given at regular intervals (L1–L22) to monitor dynamic changes in steady-state light-adapted parameters, including ΦPSII and NPQ. Following the 10-minute actinic light exposure, actinic light was turned off, and saturating pulses were applied every 30 seconds for 5 minutes to monitor the recovery kinetics of photosynthetic parameters in the dark. Second, we quantified the corrected total cell fluorescence (CTCF) of symbiotic algae by analyzing Cy5 channel images that captured their autofluorescence. The same fluorescence images used for cell counting were processed in ImageJ to calculate CTCF for individual cells, which was calculated as the integrated density minus the product of the area of the selected cell and the mean fluorescence of background readings. Autofluorescence data were collected over the 14-day cultivation period, and CTCF values were plotted using a custom python script to visualize changes in per-cell fluorescence over time.

### Protein extraction and peptide preparation for quantitative proteomics

Following the methodology previously established [54], cell lysis was conducted using a 3-in-1 extraction buffer. The resulting protein fractions, found within the bottom phase, were subsequently resolubilized in an 8 M urea buffer prepared in 25 mM Ambic (Ammonium bicarbonate). To ensure thorough and complete solubilization of the protein pellets, the mixture was vortexed and continuously agitated using a bench-top shaker at ambient room temperature. The concentration of total protein was determined spectrophotometrically utilizing the BCA Protein Assay Kit (Thermo Fisher Scientific, A55865), with bovine serum albumin (BSA) serving as the quantification standard. For preparing the proteomic sample, a quantity of 100 µg of total protein was processed via an in-solution protein extraction protocol. This procedure briefly entailed reducing the proteins with dithiothreitol (DTT) and then alkylating them using iodoacetamide (IAA). Following these modifications, enzymatic digestion was performed overnight with a high-efficiency Trypsin/Lys-C protease mix (Thermo Fisher Scientific, A40007). The resultant eluted peptides were fractionated to increase coverage using Pierce high pH reverse phase peptide fractionation columns (Thermo Fisher Scientific, 84868), and the concentration of the final peptide fractions was determined with the Pierce Quantitative Colorimetric Peptide Assay (Thermo Fisher Scientific, 23275).

### Mass spectrometry-based proteomics

Peptide analysis was performed on a timsTOF HT mass spectrometer (Bruker Daltonics GmbH) coupled to a nano Elute 2 reverse phase liquid chromatography (LC) system (Bruker Daltonics GmbH), which was equipped with a Captive Spray 2 ion source. For each analysis, 100 ng of peptide fractions from the cell samples were introduced into the nano Elute 2 LC system and separated using a 60-minute active gradient. To maintain consistent column conditions and minimize sample carryover between runs, sample injections were systematically alternated with blank injections (containing only solvent A). The specific settings for both the nano-LC system and the mass spectrometer were adopted directly from the protocol detailed in a previous study [55]. Daily monitoring of the mass spectrometer’s performance was conducted using a HeLa protein standard. Prior to initiating any sample analysis, the instrument was calibrated for both mass accuracy and ion mobility using three reference ions (*m/z* 622, 922, and 1,222) sourced from the Agilent ESI-L Tuning Mix.

### Proteomic data processing and functional annotation

Raw mass spectrometry data files (.d) were processed using FragPipe v23.1. The analysis followed the DIA_SpecLib_Quant workflow, which integrates MSFragger v4.3 [56], IonQuant v1.11.11 [57], DIA-NN v1.8.2 [58], and diaTracer v1.4.6 [59] for spectral library generation and peptide quantification [60]. Database searches were performed protein sequences predicted from *P. bursaria* CCAP 1660/18 whole-genome sequences (PRJNA659045), *P. caudatum* mitochondrial genome (NC_014262), *M. conductrix* SAG 241.80 nuclear genome (PRJNA290385), *M. conductrix* mitochondrial genome (KY629619), and *M. conductrix* chloroplast genome (NC_036806.1). Missing values were imputed according to the following stringent criteria to ensure data integrity: (1) proteins with missing values across all triplicates in a given condition were excluded from the analysis; (2) for proteins identified in only a single replicate, the observed value was propagated to the other two replicates; and (3) for proteins identified in two replicates, the missing value was substituted with the mean intensity of the existing two replicates. To estimate relative protein abundance, the intensity of each individual protein was normalized to the total sum of intensities of all identified proteins within the sample [61]. These calculated fractional abundance values were utilized for all downstream comparative analyses. Functional annotation was performed using the Kyoto Encyclopedia of Genes and Genomes (KEGG) database [62]. To ensure comprehensive coverage, orthology assignments were established by merging annotation results from both GhostKOALA [63] and KofamKOALA [64]. Subcellular localization was predicted using DeepLoc 2.1 [65]; in cases where multiple localizations were predicted for a single protein, the site with the highest probability was assigned.

## Data availability

The mass spectrometry-based proteomics data have been deposited to the ProteomeXchange Consortium via the PRIDE [66] partner repository with the dataset identifier PXD080567.

## Acknowledgements

This work was supported by grants from the NASA Exobiology Program (Grant 80NSSC24K1875) and the John Templeton Foundation (Grant 63346) to S.A.M.-G. The opinions expressed in this publication are those of the author(s) and do not necessarily reflect the views of the John Templeton Foundation. We also thank the Center for Plant Biology (CPB) at Purdue University for a seed grant that made this work possible. Electron microscopy was performed using instrumentation in the Purdue Electron Microscopy Center (RRID:SCR_022687). Samples for proteomics were analyzed at the Purdue Proteomics Facility. We gratefully acknowledge Johanna M. Dela Cruz for assistance with image analysis, Sujith Puthiyaveetil and Steven D. McKenzie for assistance with photo-physiological measurements, and Venkatesh P. Thirumalaikumar for assistance with mass spectrometry-based proteomics.

## Supplementary Figure Legends

**Figure S1.** Host cell volume along the growth curve across the nine conditions. Dots represent the mean of three biological replicates, error bars represent the standard deviation, and the solid line is a smoothed fit to the data.

**Figure S2.** Surface area-to-volume ratio (µm^-1^) of green algal photosymbionts along the growth curve across the nine conditions. Dots represent the mean of three biological replicates, error bars represent the standard deviation, and the solid line is a smoothed fit to the data.

**Figure S3.** Photosymbiont properties along the growth curve across the nine conditions. **(A)** Photosymbiont volume. **(B)** Photosymbiont number. **(C)** Symbiont-to-host volume ratio (symbiont biomass as a fraction of host volume). Dots represent the mean of three biological replicates, error bars represent the standard deviation, and the solid line is a smoothed fit to the data.

**Figure S4.** Photosynthetic properties of photosymbionts along the growth curve across the nine conditions. **(A)** Autofluorescence intensity per host. **(B)** The same data as in (A) divided by the number of symbionts per host, i.e., the value per single symbiont cell. Dots represent the mean of three biological replicates, error bars represent the standard deviation, and the solid line is a smoothed fit to the data. **(C)** Fv/Fm at Day 7. **(D)** Fv/Fm at Day 14. **(E)** NPQ at Day 7. **(F)** NPQ at Day 14. Significance of pairwise differences was assessed by t-test and is indicated above the bars (ns, not significant; *, *P* < 0.05; **, *P* < 0.01; ***, *P* < 0.001).

**Figure S5. (A)** DIC micrographs of the holobiont under the three light conditions in the high-bacteria condition; scale bars, 10 µm. **(B)** Fraction of starch within a single photosymbiont cell, measured from TEM micrographs and expressed as the percentage of single-cell volume occupied by starch. Bars represent the mean and error bars the standard deviation. Measured only in the high-bacteria condition.

**Figure S6.** Biomass-based growth curves of green algal photosymbionts across the nine conditions. **(A)** Symbiont biomass per individual host. **(B)** Symbiont biomass considering the entire host population. Dots represent the mean of three biological replicates, error bars represent the standard deviation, and the solid line is a smoothed fit to the data.

**Figure S7.** Proteome allocation in the ciliate host across the nine conditions. **(A)** Host total proteome partitioned into seven functional sectors; unlike the symbiont, the host lacks the PHO (Photosynthesis & Light Capture) sector. **(B)** Stacked bar plot of subcellular localization proteome sectors predicted with DeepLoc v2.1 [65]. In both panels, values represent the mean proteome fraction of three biological replicates.

**Table S1.** Source data (mean and SD of all plotted physiological measurements).

**Table S2.** Host proteome data (per-condition protein abundances).

**Table S3.** Symbiont proteome data (per-condition protein abundances).

**Table S4.** Proteome sector classification criteria.

**Table S5.** Functional sector summary (% of proteome per condition).

**Table S6.** Subcellular localization summary (% of proteome per condition).

**Table S7.** Symbiont plastid sub-sector summary (% of total symbiont proteome).

